# miRNAFinder: A Comprehensive Web Resource for Plant Pre-microRNA Classification

**DOI:** 10.1101/2021.06.30.450478

**Authors:** Sandali Lokuge, Shyaman Jayasundara, Puwasuru Ihalagedara, Indika Kahanda, Damayanthi Herath

## Abstract

microRNAs (miRNAs) are known as one of the small non-coding RNA molecules that control the expression of genes at the RNA level, while some operate at the DNA level. They typically range from 20–24 nucleotides in length and can be found in the plant and animal kingdoms as well as in some viruses. Computational approaches have overcome the limitations of the experimental methods and have performed well in identifying miRNAs. Compared to mature miRNAs, precursor miRNAs (pre-miRNAs) are long and have a hairpin loop structure with structural features. Therefore, most in-silico tools are implemented for pre-miRNA identification. This study presents a multilayer perceptron (MLP) based classifier implemented using 180 features under sequential, structural, and thermodynamic feature categories for plant pre-miRNA identification. This classifier has a 92% accuracy, a 94% specificity, and a 90% sensitivity. We have further tested this model with other small non-coding RNA types and obtained 78% accuracy. Furthermore, we introduce a novel dataset to train and test machine learning models, addressing the overlapping data issue in the positive training and testing datasets presented in PlantMiRNAPred for the classification of real and pseudo-plant pre-miRNAs. The new dataset and the classifier that can be used with any plant species are deployed on a web server freely accessible at http://mirnafinder.shyaman.me/.

## 1 Introduction

Ribonucleic Acid (RNA) silencing is an important and essential process that controls the class and abundance of proteins in a cell. This form of gene regulation in eukaryotes is mediated by a set of small non-coding RNAs [1]. Small non-coding RNAs are classified into four types depending on their origin, processing style, and effector protein association: microRNA (miRNA), small interfering RNA (siRNA), PIWI-interacting RNA (piRNA, animals only), and transfer RNA-derived small RNAs (tsRNAs) [2, 3, 4]. Among them, miRNAs are the most phylogenetically conserved and function post-transcriptionally to repress target genes [5, 6]. miRNAs typically range from 20–24 nucleotides in length and have been identified in plants and animals with different biogenesis. In the genome, miRNAs can be located in different regions, such as in introns, long non-coding transcripts, and/or in the chromosomal regions between two genes [7]. This study focuses on plant miRNAs and hereinafter refers to plant miRNAs as miRNAs if not otherwise specified.

The biogenesis of a miRNA starts through the transcription of miRNA genes by the RNA polymerase (Pol) II [8, 9, 10, 11]. Primary miRNA (pri-miRNA) is the initial transcript and it forms a stable hairpin-shaped stem-loop structure. Then, the pri-miRNA is processed into a pre-miRNA by a Dicer-like enzyme (DCL1) [12]. This pre-miRNA has a stem-loop hairpin secondary structure and ranges from 53nt to 938nt in length [13]. The DCL1 enzyme further processes this pre-miRNA into a mature miRNA/miRNA* duplex [14]. A small RNA methyltransferase, named Hua Enhancer (HEN1), methylates the 3’ end of the duplex [15]. Unlike in animals, the plant miRNA/miRNA* duplex is completed within the nucleus and exported into the cytoplasm by Hasty (HST), a plant homolog of Exportin-5 [16].The miRNA/miRNA* duplex is separated in the cytoplasm [17]. Loading the miRNA strand (guide strand) into the cytoplasmic Argonaute (AGO) protein (AGO1, AGO7, and AGO10 [18]) forms a ribonucleoprotein complex, RISC, to silence the target messenger RNAs (mRNAs) [17].

The miRNAs have been identified as critical regulators of developmental processes such as vegetative phase change, flowering time, leaf morphogenesis, and response adaption to environmental stresses, which are linked to most aspects of plant biology [19]. Similarly, studies have reported that plant miRNAs are involved in plant immune, hormone signaling, and signal transduction pathways [20]. Increasing evidence of the importance of miRNAs has led genetic engineering applications to a new level of designing artificial miRNAs for many potential applications to improve the agronomic properties of crops and make medicines for various human/animal/plant diseases. Therefore, the identification of existing miRNAs and their target genes is essential and a crucial task.

Studying microRNAs is challenging since microRNAs are very short, and the closely related microRNA family members differ by about one nucleotide. In the beginning, miRNAs were identified via experimental approaches such as cloning [21, 22], splinted-ligation strategy [23], and genetic screening [24]. These methods, however, are costly, time-consuming, labor-intensive, subject to chance, difficult to identify miRNAs expressed at lower levels, and difficult to clone due to physical properties such as sequence composition (Koshiol2010, Ver2009).Thus, these methods have been replaced by next-generation sequencing-based (NGS) approaches like small RNA (small RNA-Seq) sequencing in high-throughput miRNA identification [25]. The data generated by these recent technological advances requires the use of computational approaches for miRNA identification. For instance, computational systems implemented using NGS data have become the most reliable approach for large-scale miRNA detection [26, 27].

A recent review highlights the need for plant-specific computational methods for miRNA identification [28]. Based on the implementation and performance, we can classify computational methods in miRNA identification as comparative and non-comparative. Because mature miRNA sequences are short, traditional feature engineering approaches generally fail to extract effective features. As a result, most computational methods focus on identifying pre-miRNAs rather than mature miRNAs [29]. Comparative methods search for the exact or near-exact counterparts to previously validated pre-miRNAs based on the conserved characteristics of pre-miRNA sequences. Though this approach provides relative ease in detecting high-throughput evolutionarily conserved pre-miRNAs in different species, it is difficult to identify novel pre-miRNAs with distant sequence homology compared with the known pre-miRNAs [30, 31]. However, the non-comparative methods have overcome this limitation. They are based on pre-trained machine learning (ML) models that use a set of rules for pre-miRNA identification based on pre-miRNA properties such as structural, thermodynamical, and sequential variations [30, 31, 32]. Still, it requires wet-lab experiments like northern blotting or reverse transcription polymerase chain reaction (PCR) for the output validation. Further, the amount of positive and negative datasets used for the training process directly affects the performance (accuracy, specificity, and sensitivity) of the ML model [33].

Support Vector Machine [13, 34], Random forest [35], Decision trees [36] and Naïve Bayes [37] are commonly used machine learning algorithms for this task. Additionally, some studies have implemented classification models using a Stochastic-based Random Forest classifier and Naïve Bayes Classifier [38] and have been able to achieve promising results in pre-miRNA identification. According to Yousef et al. [39, 40] one-class classifiers can effectively distinguish miRNAs from other small non-coding RNA classes. Some of the studies have combined an adaptive boosting algorithm with SVM (AdaBoost-SVM) to transfer weaker classifiers into one stronger classifier [41]. This combined algorithm has shown a higher degree of efficacy compared with the other studies in mature miRNAs identification. Similarly, a limited number of studies used both homology-based methods and the ab-initio method (Machine Learning) for miRNA prediction. mirHunter [42] which identifies both plant and animal pre-miRNAs, is such an example. Neural network (NN) algorithms such as convolutional neural networks (CNN), long short-term memory neural networks (LSTM) have also been utilized in pre-miRNA identification [43, 44, 45]. Most of these models are not trained with only plant pre-miRNAs but both humans and plants pre-miRNAs. The studies have concluded that more data must be collected in order to improve identification and address the class imbalance problem. Almost all the related studies have used different versions of the miRBase [46] database as a source of a positive dataset. Most of the plant miRNA identification models have used only plant pre-miRNAs while some of them have used human, animal, virus pre-miRNAs along with plant pre-miRNAs [47, 48] for the training process. As the negative dataset, coding sequences (CDSs) are taken since miRNAs are non-coding sequences. The Phytozome database [49] and EnsemblPlants [50] are the commonly used databases for this purpose [38, 51]. However, some studies have taken other types such as non-coding small RNA sequences as the negative training and testing dataset.

The majority of studies on pre-miRNA classification have relied on data published on the web server PlantMiR-NAPred [52] [51, 53, 54, 55]. In this dataset, plant pre-miRNAs were downloaded from miRBase (version 14), which is now outdated. This study used their in-house sample selection algorithm to filter 980 sequences from each plant pre-miRNAs and pseudo-pre-miRNAs (coding sequences) to train the model. The testing dataset contains 1438 real pre-miRNAs and 1142 pseudo-pre-miRNAs. However, we discovered that there are 634 common pre-miRNA sequences among the positive training and testing sets. The model must be tested on unseen data in order to be properly evaluated. Therefore, if a study employs this dataset, it should use another positive testing dataset for the above purpose. Accordingly, we have created a new expanded gold-standard dataset derived from miRBase 22.1 for the research community.

To address the previously reported issues related to NN-based models, we developed a novel model using a multilayer perceptron (MLP) classifier, incorporating sequence-based, structural, and thermodynamic features. This model was trained and tested with our newly constructed dataset. In order to make it accessible to the research community, we deployed our classifier as a public web-server, http://mirnafinder.shyaman.me/. Similarly, the developed dataset is also available on the same server. This article reports the dataset generation, feature calculation, model training, result evaluation, and deployment of the classifier.

## 2 Methodology

### 2.1 Data preparation

In this study, we present a new dataset to the research community. This section discusses the criteria we followed to construct the complete dataset to train and test the pre-miRNA classification models.

#### 2.1.1 Positive dataset

We downloaded plant pre-miRNAs from *miRBase* [46] version 22.1, which serves as the primary online repository for miRNA sequences and annotations. This database comprises 38,589 hairpin precursor miRNA entries, which express 48,885 mature miRNAs from 271 species. We retrieved 8615 plant pre-miRNA sequences and eliminated 261 duplicates. Then we discarded 106 sequences that contained nucleotides other than Adenine (A), Guanine (G), Cytosine (C), and Uracil (U). After the initial preprocessing stage, we obtained a total of 8,248 pre-miRNAs.

To generate a generalised test set, we filtered out pre-miRNA records that did not have at least one experimentally validated mature miRNA for the pre-miRNA and at least two referenced articles. The final positive training dataset consisted of 1,211 pre-miRNAs after meeting the above criteria. Figure 1 illustrates the length distribution of the 1,211 real pre-miRNAs. This set includes pre-miRNAs from *Arabidopsis lyrata, Arabidopsis thaliana, Brachypodium distachyon, Brassica napus, Brassica rapa, Chlamydomonas reinhardtii, Citrus sinensis, Gossypium hirsutum, Glycine max, Hordeum vulgare, Malus domestica, Manihot esculenta, Medicago truncatula, Nicotiana tabacum, Oryza sativa, Picea abies, Prunus persica, Physcomitrella patens, Pinus taeda, Populus trichocarpa, Sorghum bicolor, Solanum lycopersicum, Solanum tuberosum, Vitis vinifera, and Zea mays*.

**Figure 1:**
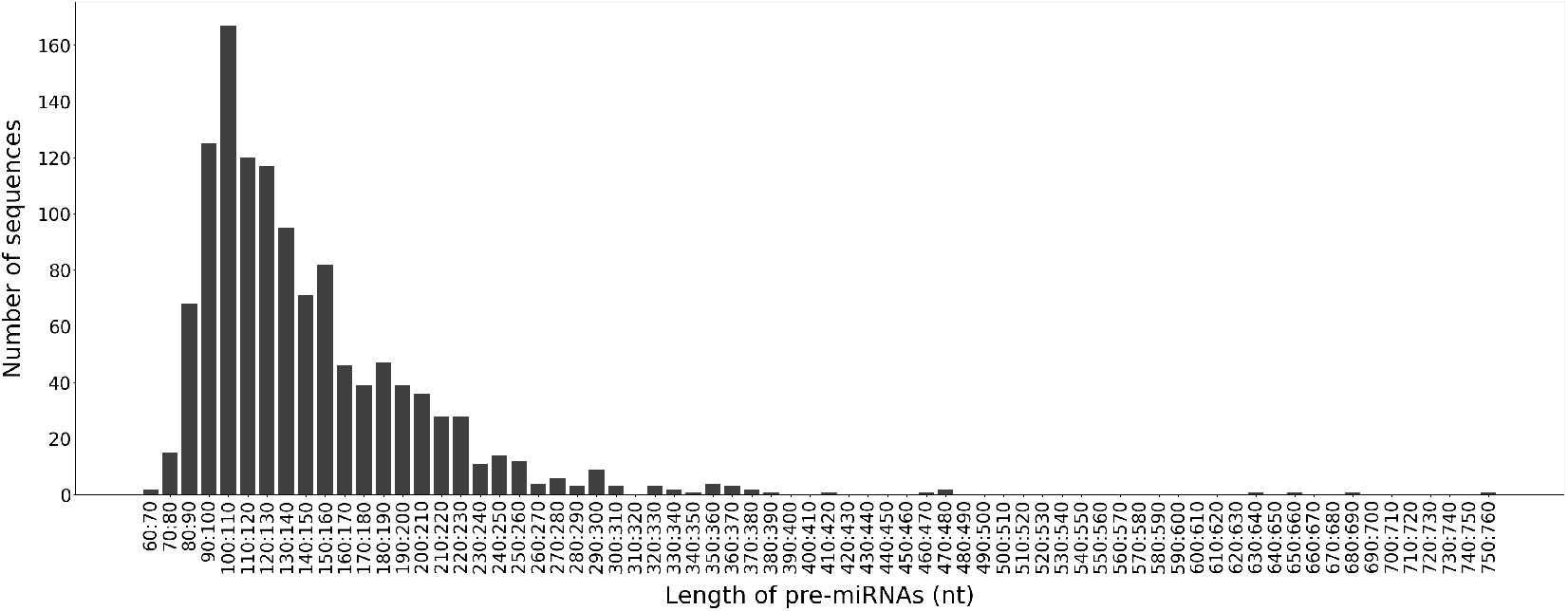
Length distribution of pre-miRNA sequences in training data

The remaining 7037 pre-miRNAs were classified into species and ranked according to sequence count. The testing dataset then included 100 sequences from each of the top ten ranked species. *Brachypodium distachyon* (bdi), *Glycine max* (gma), *Gossypium raimondii* (gra), *Lotus japonicus* (lja), *Medicago truncatula* (mtr), *Oryza sativa* (osa), *Picea abies* (pab), *Populus trichocarpa* (ptc), *Sorghum bicolor* (sbi), and *Solanum tuberosum* (stu) were among the selected species.

#### 2.1.2 Negative dataset

To create the negative dataset, coding DNA sequences (CDS) were used as the pseudo pre-miRNA hairpin structure sequences. The EnsemblPlants (Release 47) database was used to obtain CDS of *Arabidopsis thaliana*, *Triticum aestivum*, *Oryza sativa*, *Hordeum vulgare*, *Zea mays*, and *Physcomitrella patens*. We chose CDS only in the range of 70nt to 260nt to be comparable with the positive training set (Figure 1). To be effective, the model should be able to differentiate between positive and negative sequences with similar GC content, as well as sequences with similar Minimum Free Energy (MFE). Accordingly, we created a negative training dataset that was similar to the positive training dataset by employing the G+C% and MFE as described below.

Since testing and training datasets have to be mutually exclusive, we first removed 1000 pre-miRNA sequences in the testing dataset from the preprocessed 8248 pre-miRNAs. The MFE and G+C% of the resulting 7248 plant pre-miRNAs and CDSs were then calculated using RNAfold from the Vienna package [56] with the default parameters and grouped according to their lengths. Then, for each length group, we calculated the minimum and maximum values of MFE and G+C percent. The CDSs that meet the following two criteria were then extracted from each group.

- The MFE of a CDS in a particular length range should be between the minimum and maximum MFE values of the positive sequences in the same length range **AND**
- The G+C% of the same CDS should be between the minimum and maximum G+C% of the positive sequences in the same length range

For instance, a coding sequence in length from 70 to 79 nt should satisfy 9.6 <= MFE <= 70.9 and 17.16 <= G+C % <= 75.71.

The CDSs filtered using the aforementioned criteria were then randomly sampled in the same proportion as in the length intervals of the positive training set. Assume that the ratios for the length groups 70–79, 80–89, 90–99, 100–109, and 110–119 of the positive training sequences are 0.013, 0.059, 0.107, 0.144, and 0.103. The negative training samples should then have the same proportion for each length group as the positive training samples. Finally, we obtained 1,211 CDSs as the negative training dataset. The length distribution of the 1,211 CDSs in the negative training dataset is presented in the Figure 2.

**Figure 2:**
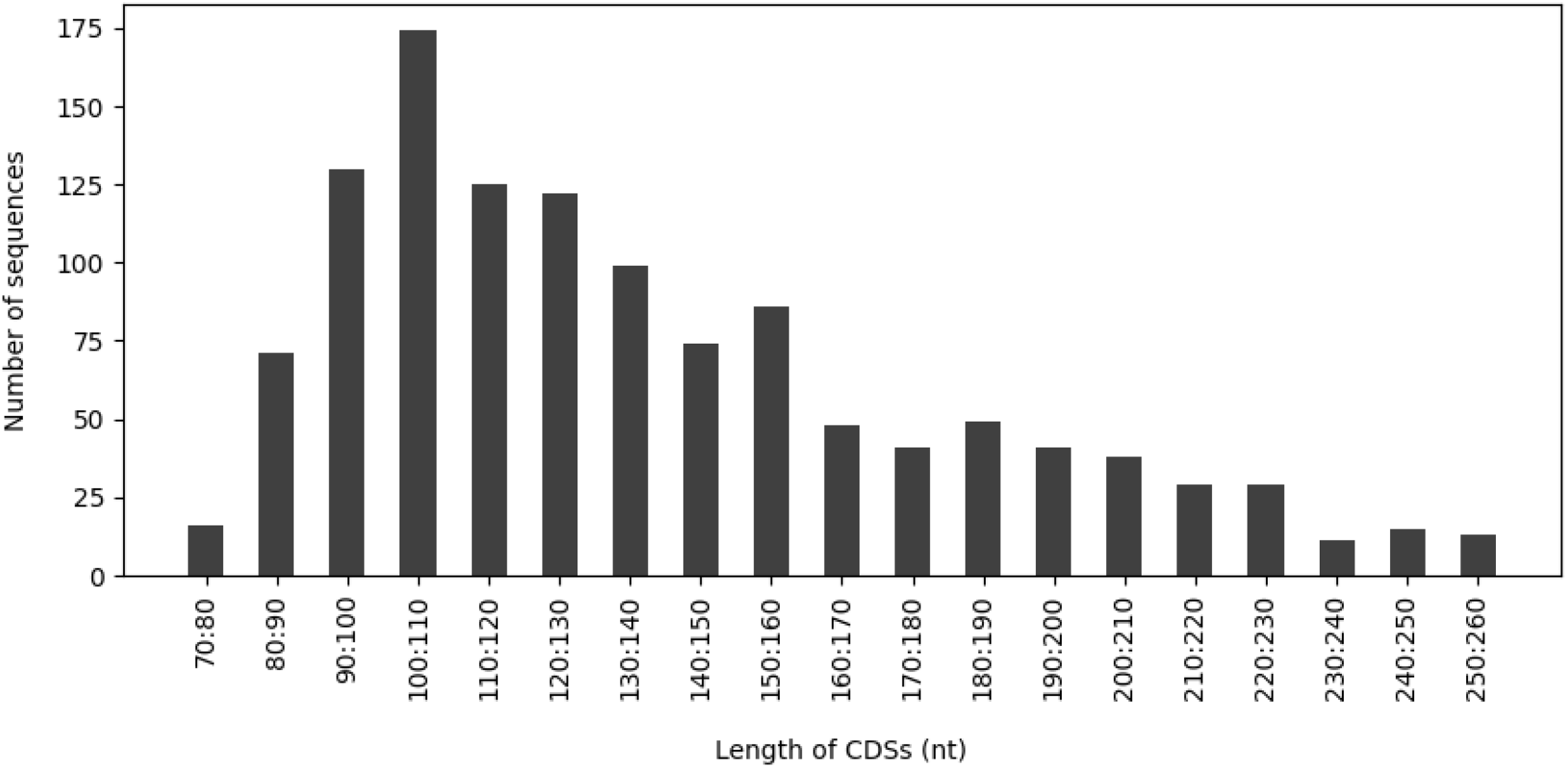
Length distribution of negative (CDS) training sequences

We used the same negative testing dataset in PlantMiRNAPred [52] because there were no shared sequences with our negative training dataset. This negative testing dataset was comprised of 1,142 CDSs. In addition to this testing dataset, we decided to collect additional non-coding RNA sequences to see how the model differentiates pre-miRNA sequences from other non-coding RNAs. A robust predictive model should be able to distinguish pre-miRNA not only from CDSs but also from other non-coding sequences. We downloaded other non-coding RNA sequences from RNAcentral (v15) [57], a non-coding RNA sequence database. Using the following search query, we included only sequence records that had been verified by at least three databases and had a secondary structure. The downloaded collection included 43 sequences of ribosomal RNA (rRNA), 579 sequences of small nucleolar RNA (snoRNA), 431 sequences of small nuclear RNA (snRNA), and 1,706 sequences of transfer RNA (tRNA).

- plant* AND expert_db: “Ensembl Plants” AND expert_db: “Rfam” AND expert_db: “ENA” AND has_secondary_structure: “True” AND rna_type: “rRNA”

Accordingly, our negative testing dataset included 1,142 CDSs and 2,759 other non-coding RNA sequences.

### 2.2 Features

Recent studies have introduced a set of features in both primary sequence and secondary structure that can be used to distinguish real plant pre-miRNAs and pseudo hairpins [52, 58, 35, 53]. We used 180 features under sequential, structural, and thermodynamic feature categories. We used scripts and algorithms described in our previous study, [59] to extract the features. Accordingly, this study includes 48 features introduced in microPred [60], triplet element features, and motif features. Our previous research [59], however, concluded that motif sequences have less of an impact on pre-miRNA identification than the other features. Furthermore, taking the computational time required for motif discovery into account, we extracted only 50 motifs with more than 10 sites from each positive and negative training set.

We generated the correlation matrix to get an overview of the features and discovered that there are certain highly correlated variables among them. Then we perform dimensionality reduction using *Principal Component Analysis (PCA)* [61]. Before performing the *PCA*, we normalized all the variables. A cumulative variance graph was plotted against the number of components, using the explained variances to determine the number of principal components. We selected 123 as the number of components where the cumulative variance reached 97% (Figure 3).

**Figure 3:**
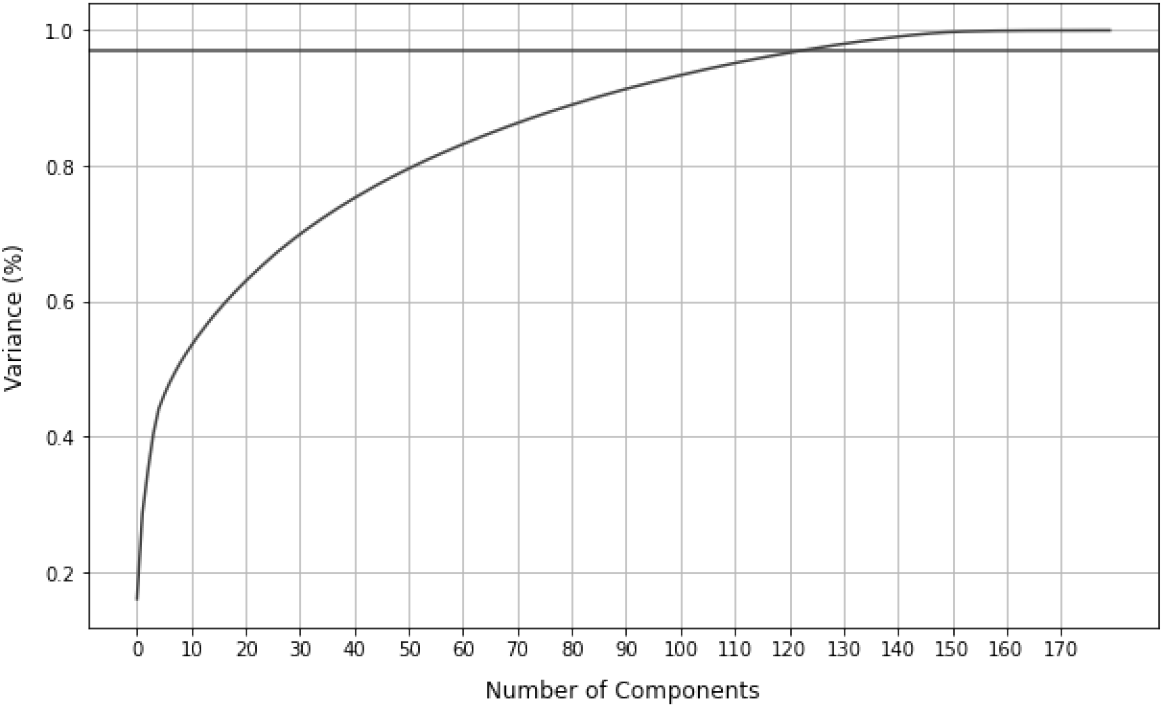
Cumulative variance vs number of principal components

### 2.3 Model training and performance evaluation

The past literature on miRNA identification has largely employed classification methods such as SVM, Decision tree, Random Forests, and Naive Bayes to develop their models. However, relatively few studies have focused on deep learning and neural network-based techniques for plant miRNA classification because of the limitation on obtaining a large number of positive miRNAs to train the model. Since we have 2,422 sequences for training, we employed a Multi-Layer Perceptron Classifier (MLP), which relies on an underlying neural network to execute the task of classification. After executing the hyperparameter tuning, the MLP model was trained with the Relu activation function, a constant learning rate, and 3 hidden layers with (100, 50, 20) units each.

To evaluate the model’s performance, we use four criteria: accuracy, specificity, sensitivity, and area under the receiver operating characteristics curve (AUC-ROC).

## 3 Results and Discussion

### 3.1 Model evaluation

The model was developed using 5-fold cross-validation on 2422 sequences. The training accuracy at 5-fold crossvalidation was 98.8% with a standard deviation of 0.005. The model was evaluated for classification accuracy on a test set of 1000 plant pre-miRNA sequences and 1142 CDSs. Accuracy, specificity, and sensitivity were 92%, 94%, and 90% respectively. Testing the model with actual pre-miRNAs and CDSs gives a 0.92 AUC-ROC value, indicating the high capability of differentiating the pre-miRNA sequences from coding sequences.

We further tested the model with actual pre-miRNAs and other non-coding RNAs and obtained an AUC-ROC value of 0.84. This result indicates that the developed model has a considerable class separation capacity for distinguishing pre-miRNAs from other non-coding RNA sequences. We obtained 78% testing accuracy for other non-coding RNA types. Though the negative training dataset doesn’t include sequences from these non-coding RNA types, the model has performed well for such sequences as well. Furthermore, we tested our model for each non-coding RNA type individually. We obtained 70%, 72%, 90%, and 76% accuracy values for rRNA, snRNA, snoRNA, and tRNA, respectively.

We checked for the species-wise accuracies (Table 1) on the positive test set. All the species have an accuracy of greater than 75%. However, some species like *Brachypodium distachyon* (bdi), *Medicago truncatula* (mtr) and *Oryza sativa* (osa) have fewer accuracies compared to other species.

**Table 1:** Testing accuracy for different species

| Dataset | MLP model accuracy |
| --- | --- |
| Brachypodium distachyon | 0.82 |
| Glycine max | 0.90 |
| Gossypium raimondii | 0.94 |
| Lotus japonicus | 0.96 |
| Medicago truncatula | 0.79 |
| Oryza sativa | 0.82 |
| Picea abies | 0.91 |
| Populus trichocarpa | 0.93 |
| Sorghum bicolor | 0.95 |
| Solanum tuberosum | 0.95 |

Comparing our results with past studies is not reasonable as we employed a novel training and testing dataset. However, a few research have utilized a neural network-based classification model for plant pre-miRNA identification. We have acquired significant accuracy, specificity, sensitivity, and AUC-ROC for plant pre-miRNA identification.

We also examined the overall (Supplementary table S1) and species-specific (Supplementary table S2) classification performance of the models trained on each feature category (i.e., Micropred features, triplet element features, and motif features). Models were trained on both PCA and non-PCA features. In addition, we used chi-square scores to rank the original features (without PCA) to find the relationship between the features and the target variable (Supplementary table S3).

### 3.2 Comparison of the dataset

We compared our positive training and testing datasets to that of PlantMiRNAPred. PlantMiRNAPred consists of 980 sequences in the training set and 1438 sequences in the testing set. However, 634 common sequences belong to both the training and testing sets. Thus, the PlantMiRNAPred dataset had 1784 sequences in total. The dataset in this study comprises of 1211 sequences in the training dataset and 1000 sequences in the testing dataset, with no overlapping sequences.

We used sequence ID as the sequence compactor and found that there were 801 common sequences in both datasets (Figure 4). But miRNAFinder has 1410 new sequences since it is derived from miRBase v22.1.

**Figure 4:**
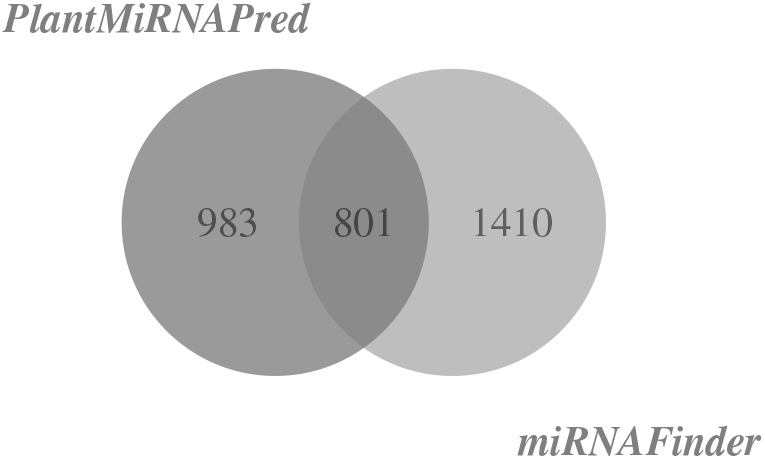
Comparison of PlantMiRNAPred dataset with miRNAFinder dataset

In addition, we compared the pre-miRNA sequences in the training datasets of the two studies. There were 304 sequences that were shared by both training sets, 907 novel sequences in miRNAFinder, and 676 sequences that were unique to the PlantMiRNAPred dataset. When the two testing datasets were compared, there were 190 similar sequences. Similarly, 810 and 1248 distinct sequences were found solely in miRNAFinder and PlantMiRNAPred, respectively. The low number of overlapping sequences in the testing datasets could be due to the fact that miRNAFinder contains pre-miRNAs of plant species such as *Brachypodium distachyon*, *Gossypium raimondii*, *Lotus japonicus*, *Picea abies*, *Sorghum bicolor* and *Solanum tuberosum* that have not been considered in previous studies.

## 4 miRNAFinder web server

We published our classifier, miRNAFinder, on a web server that anyone could use to predict potential miRNAs. The input sequences should be in FASTA format, and there are two ways to enter them. The user can either enter the FASTA sequences into the provided text box or upload them using the “Choose file” button (Figure 5). We set the maximum number of sequences to 10 because feature calculation is computationally intensive and takes a long time. For each sequence, the output table comprises the sequence ID, predictions, and confidence scores (Figure 6). The results and extracted features are available for download from the same page. The training and testing datasets can also be downloaded from the resources page.

**Figure 5:**
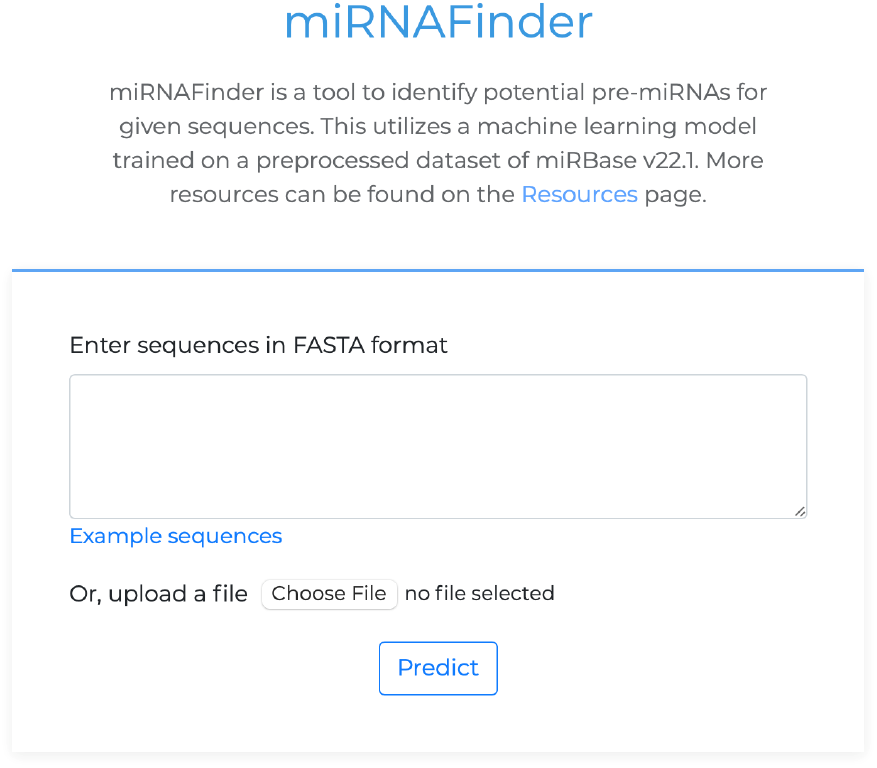
Homepage of the miRNAFinder web server

**Figure 6:**
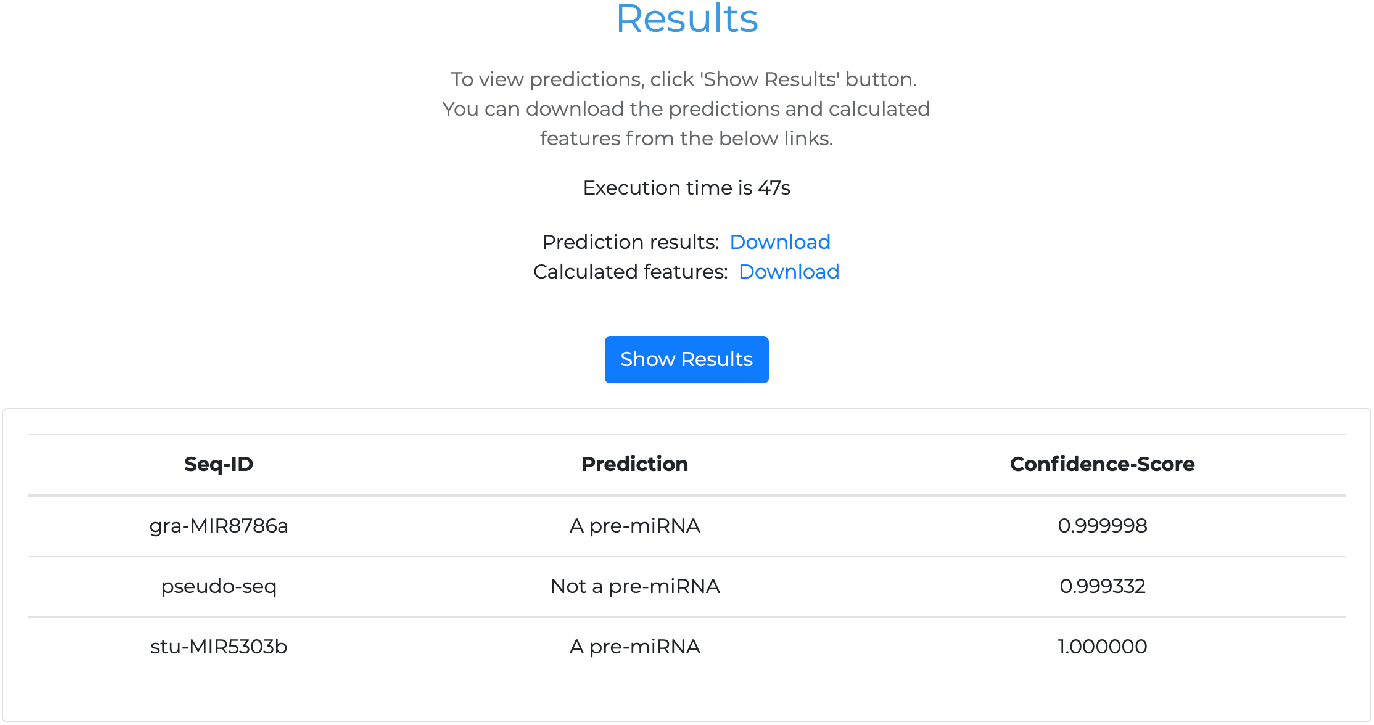
Results view of the miRNAFinder web server

**Figure 7:**
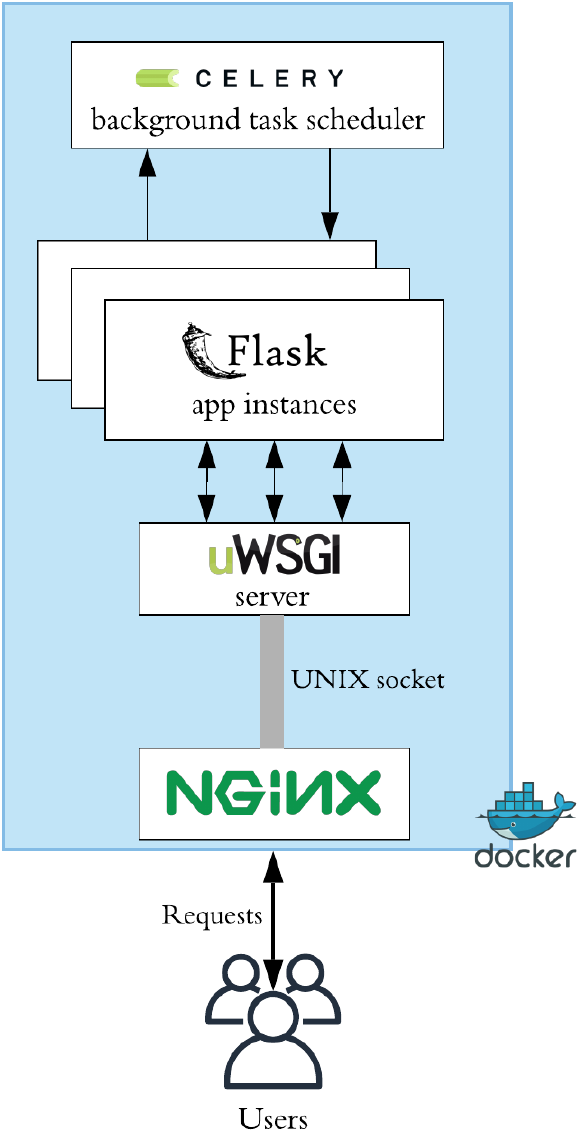
miRNAFinder architecture

Flask, a Python web framework, was used to create the miRNAFinder web app. Multiple instances of the miRNAFinder flask program were run on the uWSGI application server, and user requests were handled efficiently. We used Celery, a background job scheduler, to improve server availability because feature extraction is a time-consuming task. Nginx functions as a reverse proxy as well as a load balancer. To allow for easier deployments, the entire program was built within Docker containers. Figure fig:architecture depicts the whole application architecture of miRNAFinder.

## 5 Web tool and data availability

miRNAFinder can be found at http://mirnafinder.shyaman.me/. The source code for the miRNAFinder server is publicly available on Github (https://github.com/shyaman/miRNAFinder-web-server) and in the Zenodo archive (https://doi.org/10.5281/zenodo.4719605). Anyone can set up their own instance of the miRNAFinder server based on their needs (for example, when a user needs to make predictions for more than ten sequences). The high-quality dataset presented here is available at https://doi.org/10.5281/zenodo.4721396.

## 6 Conclusion and future work

Classification of real and pseudo-pre-miRNAs in plant species is challenging since plant pre-miRNAs have a diverse range of lengths compared to the conserved length of animal pre-miRNAs. For the scientific community, we present a new dataset of real and pseudo-pre-miRNAs. It includes 2211 real pre-miRNAs from miRBase (v22.1), 2353 CDSs from the EnsemblPlants database, and 2759 additional non-coding RNAs from the RNAcentral database. The scripts in the *microPred* were used to calculate compositional and thermodynamic features. Despite the fact that *microPred* concentrated on human miRNAs, the features had no detrimental impact on performance. The MLP classifier categorized pre-miRNAs with 92% accuracy, 94% specificity, and 90% sensitivity. The results suggest that the MLP classifier combined with the mentioned features can achieve significant accuracy and an AUC-ROC value. In future work, we suggest assessing the performance of MLP against classifiers such as RF, NB, and SVM using the new dataset.

In terms of the computational time, calculating the features introduced in the *microPred* took the longest time. In this work, the number of sequences used in training was reduced based on multiple criteria to reduce the computational time. It will be worthwhile to investigate alternate ways of reducing computing time for feature extraction without compromising the amount of data utilized for training.

The proposed model and dataset are accessible through a user-friendly web-server, *“miRNAFinder”*. The freely available dataset indicated above can be used to create, implement, and test new methods for plant pre-miRNA prediction and can be used with any species.

