## Supplementary for "miRNAFinder: A Comprehensive Web Resource for Plant Pre-microRNA Classification"

### Supplementary Tables

**Table S1: Comparison of the models trained with different feature sets on all test data**

PCA - Principal component analysis
Micropred – Features taken from the study, microPred (DOI: 10.1093/bioinformatics/btp107)
Motif – Motif features
Triplet – Triplet element features

| **Feature set** | **With PCA** | | | **Without PCA** | | |
|---|---|---|---|---|---|---|
| | **Accuracy** | **Sensitivity** | **Specificity** | **Accuracy** | **Sensitivity** | **Specificity** |
| All | 0.87 | 1 | 0.72 | 89 | 100 | 76 |
| Micropred | 0.92 | 0.98 | 0.84 | 92 | 99 | 84 |
| Motif | 0.62 | 0.49 | 0.77 | 70 | 73 | 67 |
| Triplet | 0.84 | 0.95 | 0.71 | 82 | 97 | 66 |

**Table S2: Accuracy comparison of the models trained with different feature types on the ten different plant species**

PCA - Principal component analysis
Micropred – Features taken from the study, microPred (DOI: 10.1093/bioinformatics/btp107)
Motif – Motif features
Triplet – Triplet element features

| **Plant species** | **Without PCA** | | | | **With PCA** | | | |
|---|---|---|---|---|---|---|---|---|
| | **Micropred** | **Motif** | **Triplet** | **All** | **Micropred** | **Motif** | **Triplet** | **All** |
| Brachypodium distachyon | 0.84 | 0.84 | 0.19 | 0.79 | 0.77 | 0.87 | 0.22 | 0.89 |
| Glycine max | 0.91 | 0.83 | 0.39 | 0.81 | 0.83 | 0.87 | 0.41 | 0.86 |
| Gossypium raimondii | 0.93 | 0.98 | 0.13 | 0.85 | 0.84 | 0.95 | 0.17 | 0.94 |
| Lotus japonicus | 0.85 | 0.94 | 0.28 | 0.77 | 0.71 | 0.88 | 0.27 | 0.85 |
| Medicago truncatula | 0.79 | 0.96 | 0.31 | 0.69 | 0.72 | 0.91 | 0.35 | 0.81 |
| Oryza sativa | 0.83 | 0.92 | 0.18 | 0.72 | 0.79 | 0.87 | 0.25 | 0.84 |
| Picea abies | 0.93 | 0.82 | 0.21 | 0.87 | 0.87 | 0.75 | 0.24 | 0.92 |
| Populus trichocarpa | 0.90 | 0.84 | 0.29 | 0.84 | 0.87 | 0.77 | 0.32 | 0.87 |
| Sorghum bicolor | 0.95 | 0.72 | 0.12 | 0.66 | 0.89 | 0.75 | 0.13 | 0.89 |
| Solanum tuberosum | 0.88 | 0.82 | 0.35 | 0.90 | 0.80 | 0.93 | 0.37 | 0.90 |

**Table S3: Top 50 features with the highest chi-square score**

| **Rank** | **Feature** | **Description** | **Score** |
|---|---|---|---|
| 1 | dQ | Normalized shannon entropy | 177.2054581 |
| 2 | dD | Adjusted base pair distance | 172.3157993 |
| 3 | zQ | Normalized variant of dQ | 114.6060978 |
| 4 | zD | Normalized variant of dD | 112.5106023 |
| 5 | Tm | Melting Energy of the structure | 92.52089757 |
| 6 | diff | \|MFE - ensemble free energy\|/ Length | 87.97627547 |

| | | | |
|---|---|---|---|
| 7 | dP | Normalized base-pairing propensity | 86.80169311 |
| 8 | A(.. | Triplet element | 79.49092067 |
| 9 | zF | Normalized variant of dF (The second (Fielder) eigenvalue) | 70.86020584 |
| 10 | zP | Normalized variant of dP (Normalized base-pairing propensity) | 65.70636001 |
| 11 | auL | Normalized base pair count | 64.31542141 |
| 12 | mfe1 | dG / %(C+G) | 62.76117561 |
| 13 | A..( | Triplet element | 59.62084541 |
| 14 | freq | Frequency of MFE (Minimum Free Energy) structure | 56.97380484 |
| 15 | A... | Triplet element | 52.29025838 |
| 16 | bpStems | Average base pairs per stem | 45.30330139 |
| 17 | auStems | %(A-U)/number of stems | 43.77481921 |
| 18 | dG | Normalized minimum free energy of folding | 42.29900855 |
| 19 | C((. | Triplet element | 40.09264109 |
| 20 | G.(. | Triplet element | 39.90636973 |
| 21 | G..( | Triplet element | 39.43294806 |
| 22 | div | Divergence between positive and negative classes | 38.68833766 |
| 23 | C.(. | Triplet element | 37.01158928 |
| 24 | nefe | Normalized Ensemble Free Energy | 36.79277189 |
| 25 | dF | The second (Fielder) eigenvalue | 36.58590572 |
| 26 | G(.. | Triplet element | 35.28312445 |
| 27 | mfe4 | MFE index 4 (MFE/total bases) | 32.04877891 |
| 28 | dHL | Normalized Structure Enthalpy | 31.52881541 |
| 29 | G((. | Triplet element | 28.97650792 |
| 30 | dSL | Normalized Structure Entropy | 28.74267168 |
| 31 | zG | Normalized variant of dG | 28.65685095 |
| 32 | A(.( | Triplet element | 28.50996605 |
| 33 | C... | Triplet element | 27.66351448 |
| 34 | C.(( | Triplet element | 27.10584775 |
| 35 | %CG | CG dinucleotide frequency | 27.06810536 |
| 36 | A((( | Triplet element | 22.9811094 |
| 37 | C..( | Triplet element | 22.96685613 |
| 38 | G(.( | Triplet element | 21.87127775 |
| 39 | G... | Triplet element | 20.95645354 |
| 40 | U.(. | Triplet element | 20.8547232 |
| 41 | motif50 | Motif DBCAVRMCAUGHMKGYSCHUWSUUCUYCWAYYYVASWA | 18.97558822 |
| 42 | C(.. | Triplet element | 16.16861024 |
| 43 | G.(( | Triplet element | 14.28973841 |
| 44 | motif2 | Motif YUYUKUKNUUKGAUUGARSSGWGCYMMUM | 14.23691869 |
| 45 | %UU | UU dinucleotide frequency | 13.79627474 |

| | | | |
|---|---|---|---|
| 46 | gcStems | %(G+C)/number of stems | 13.49945208 |
| 47 | mfe3 | MFE index 3 (dG/no. of loops in secondary structure) | 12.42992395 |
| 48 | motif44 | Motif UYYYUCUYUYUCUCU | 11.20947997 |
| 49 | motif0 | Motif UCRGMCMAGGMUKMMUUSCCS | 10.52319426 |
| 50 | U.(( | Triplet element | 10.42463588 |
